# An upstream open reading frame represses translation of the neuronal potassium channel KCNQ2

**DOI:** 10.64898/2026.03.24.713967

**Authors:** Dalton J. Huey, Eduardo Guadarrama, Jean-Marc DeKeyser, Carlos G. Vanoye, Christine Q. Simmons, Qianru Li, Emily K. Stroup, Zhe Ji, Alfred L. George

## Abstract

Upstream open reading frames (uORFs) within the 5’-untranslated region (5’-UTR) of mRNA transcripts can regulate protein translation. Despite widespread prevalence within the human genome, they remain unidentified for many clinically relevant genes. A gene frequently associated with neonatal-onset epilepsy is *KCNQ2*, which encodes a neuronal voltage-gated potassium channel subunit that functions to dampen neuronal excitability. Heterozygous loss-of-function pathogenic *KCNQ2* variants are known to cause a range of neurodevelopmental disorders and epileptic encephalopathies, but there remains an unmet clinical need for patients harboring these variants. We identified a single uORF in *KCNQ2* that is highly repressive of protein translation and demonstrated that mutations disabling the uORF start codon enhance translation of encoded potassium channels. Additionally, we show that adenine base editing of the uORF start codon can weaken ribosome engagement at the uORF and enhance translation of the protein in a neuron-like cell line. This study establishes a previously underexplored regulatory feature for *KCNQ2* and highlights the importance of understanding uORFs for clinically relevant genes, both for assessing disease risk and therapeutic potential.

## Introduction

The 5’-untranslated region (5’-UTR) of mature mRNA transcripts often contain regulatory features that govern the rate of protein translation (1). Despite emerging knowledge about the function of these features, noncoding regions remain underexplored for many clinically relevant genes (2). In particular, upstream open reading frames (uORFs) are present in approximately 50% of coding genes in the human transcriptome, of which 20% are estimated to actively regulate translation (3, 4). These consist of ATG or other near-cognate start codons within the 5’-UTR that produce short open reading frames (ORFs) either terminating before, or overlapping with, the canonical start codon in a different reading frame (5). Generally, uORFs repress translation of the main ORF by sequestering 43S ribosome preinitiation complexes to the upstream start codon during the scanning process before reaching the canonical start codon (6). Translation of the canonical ORF for uORF-containing genes involves scanning past the upstream start codon (leaky scanning) or through reinitiation after translating the uORF, although the degree to which these events occur can vary (7–10).

Several approaches have demonstrated that disabling uORF start codons can enhance translation of the canonical ORF, establishing uORFs as therapeutic targets for monogenic disorders of haploinsufficiency. This was shown using steric-blocking antisense oligonucleotides (ASOs) targeting the uORF start codon, (11, 12), although the reproducibility of this approach and its relevance in certain gene systems have been debated (13–15). Newer approaches have achieved a similar effect through direct DNA or RNA base editing to convert the uORF start codon to a translationally inefficient codon (16, 17), or indirectly through ASO-mediated modulation of RNA structure to weaken uORF translation (18). For genes containing uORFs in a different exon from the canonical start codon, exon skipping has also been used as a strategy to relieve uORF repression on target genes (16, 19, 20). Aside from the therapeutic potential, uORFs are also an underexplored gene element potentially involved in disease pathogenesis. For example, mutations disrupting the uORF stop codon or changing the reading frame can extend existing uORFs, resulting in repression of canonical ORF translation beyond physiological levels (21–23). Identification of uORF elements in clinically relevant genes can thus be informative both as a therapeutic target and as a potential disease mechanism.

One example of a gene with unmet clinical need is *KCNQ2*. Heterozygous loss-of-function variants in *KCNQ2* cause a spectrum of monogenic neurodevelopmental disorders with epilepsy (24–26). The severity of the condition can range from self-limited familial neonatal epilepsy (SLFNE), where seizures resolve over time (27), to severe developmental and epileptic encephalopathy (DEE), which can present with motor defects, intellectual disability, and seizures persisting throughout life (28, 29). Currently, there is no marketed drug available for these diseases through a *KCNQ2*-targeted approach.

The *KCNQ2* gene encodes a voltage-dependent potassium channel (KCNQ2 or K_V_7.2) that is expressed throughout the brain (30, 31). Functional channels are produced as homotetramers of four KCNQ2 subunits, or more commonly as heterotetramers with KCNQ3 (K_V_7.3), with the latter conducting larger potassium currents (32, 33). The potassium conductance resulting from the opening of these channels constitutes the M-current, a slow activating, non-inactivating current that opens at subthreshold potentials to oppose excitatory stimuli (34). As a result, cells with dysfunctional channels are hyperexcitable and unable to dampen repetitive action potential firing, which are hallmarks of seizures and epilepsy (35).

In this study, we investigated the *KCNQ2* 5’-UTR using bioinformatic and experimental approaches to search for functional uORF sequences. We discovered that a single uORF in *KCNQ2* is a strong translational repressor, and that mutations disrupting the uORF start codon promote higher levels of KCNQ2 protein.

We further demonstrated that adenine base editing of the upstream start codon in a neuron-like cell line disables the uORF and boosts KCNQ2 protein expression. Our work implicates a functional uORF as a translational regulator of the epilepsy gene *KCNQ2* and suggests a potential target for modulating expression of this important protein.

## Results

### The *KCNQ2* 5’-UTR contains an actively translated uORF

We identified a single ATG start codon (AUG in RNA, all subsequent mentions will be written with the DNA notation) within the human *KCNQ2* 5’-UTR that precedes a putative 66 nucleotide (nt) uORF terminating 37 nt prior to the canonical start codon (**Fig. 1A**). The uORF start codon is within a moderate Kozak context, containing a purine at the −3 position, but lacking a guanine at the +4 position. The uORF is GC-rich (88% GC content) and its predicted encoded peptide sequence is largely composed of proline and glycine residues. Other near-cognate start codons including GTG and CTG were also present within this sequence, however, we focused our attention on the sole ATG codon, which typically initiate stronger translational repression compared to near-cognate start codons.

**Fig. 1:**
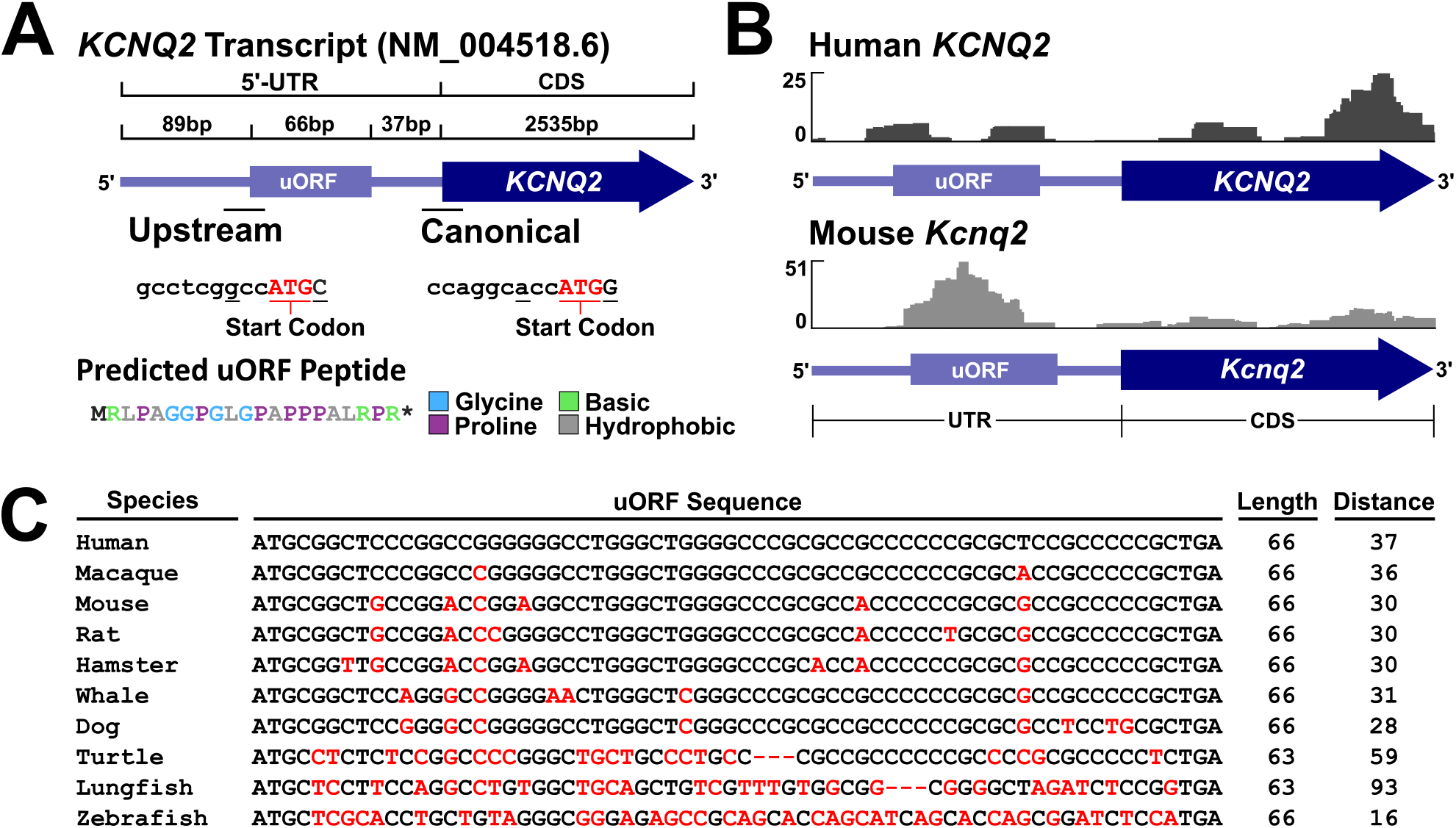
The *KCNQ2* 5’-UTR contains a conserved and actively translated uORF sequence. **(A)** Schematic of the human *KCNQ2* 5’-UTR sequence. The position of the uORF within the 5’-UTR is shown, relative to the coding sequence (CDS) of the *KCNQ2* gene. The Kozak consensus sequence is listed for the upstream and canonical start sites, with the start codon annotated in red and the −3 and +1 nucleotides of the motif underlined. The predicted peptide sequence from translation of the uORF is displayed (glycine, blue; proline, purple; basic, green; hydrophobic, gray). **(B)** Ribosome profiling of published datasets from human and mouse brain. Mapped reads of ribosome-protected footprints are shown for the uORF and the 5’-proximal *KCNQ2* CDS. Reads are combined from five individuals for the human dataset, and three independent mice for the mouse dataset. **(C)** Sequence alignment of the *KCNQ2* uORF sequence across species by Clustal Omega. Mismatches and deletions in relation to the human sequence are shown in red. The uORF length and distance from the stop codon to the canonical start codon for each sequence is also listed.

To determine whether the putative *KCNQ2* uORF undergoes active translation, we examined ribosome profiling datasets obtained from human and mouse brain (36). Specifically, these datasets included human brain tissue from surgical resections for glioma or non-neoplastic tissue for treating epilepsy, while mouse brain samples were from normal frontal lobe tissue. Ribosome-protected reads mapped to the *KCNQ2* uORF in both species, indicative of active translation (**Fig. 1B**). The uORF sequence has a high degree of nucleotide sequence identity between human *KCNQ2* and mouse *Kcnq2* orthologs. Consistent with these findings, similar results were observed using the GWIPS-viz online browser of ribosome profiling data for both human and mouse transcriptomes (37).

### Evolutionary conservation of the *KCNQ2* uORF

To further evaluate the conservation of the uORF, we collected sequences of *KCNQ2* orthologs from the NCBI database, with a particular focus on model organisms and including at least one representative species from each major vertebrate clade. We examined the sequence just upstream of the canonical start codon to identify uORFs of a similar length containing an ATG start codon and conducted a multiple sequence alignment.

Strong conservation was observed among mammals, with each uORF sequence sharing the same 66 nt length starting with an ATG start codon and terminating with a TGA stop codon (**Fig. 1C**). We did not observe similar sequences in multiple non-mammalian species including birds, alligators, lizards, and rays, but the more distantly related turtle and lungfish had a uORF sequence that was 3 nt shorter than mammalian *KCNQ2* uORF, retaining the same reading frame. These findings suggest there is evolutionary pressure to maintain this feature in *KCNQ2*.

### *KCNQ2* uORF is a strong repressor of canonical translation

We determined the extent to which the *KCNQ2* uORF influences translation of the canonical ORF using reporter gene assays. Given the high degree of conservation between human and mouse uORF sequences, we compared reporter gene translation under control of the 5’-UTR from each species. We inserted the human and mouse *KCNQ2* 5’-UTR sequences immediately preceding the coding region of nanoluciferase (NanoLuc) in a mammalian expression plasmid (**Fig. 2A)**. Compared to the unmodified *Nluc* construct, insertion of the *KCNQ2* 5’-UTR sequence repressed reporter gene translation by 7-fold for the human sequence (p=2.26E-9) and 3-fold for the mouse sequence (p=1.73E-7) (**Fig. 2B**). We then introduced single nucleotide mutations of the uORF ATG start codon to create an AAG near-cognate start codon, which was previously reported as one of the most inefficient at initiating translation (38). Upon introducing the AAG uORF mutations, the mutant human 5’-UTR exhibited a 5-fold higher level of reporter gene translation compared with the wildtype sequence (p=2.41E-6), whereas the mutant mouse 5’-UTR showed a 2-fold higher level (p=1.35E-5). These findings indicate that the upstream start codon is responsible for a large portion of the repressive effect within the *KCNQ2* 5’-UTR for both human and mouse sequences. While the fold-change for the mouse 5’-UTR appears low, this effect appears to be primarily driven by less overall repression of the 5’-UTR sequence rather than a weaker uORF. Absolute differences in firefly-normalized luminescence show a similar degree of uORF-dependent repression between human and mouse sequences (human: 19.3, mouse: 17.1, p=0.57 for difference using Welch’s t-test), suggesting that the degree of repression on translation is similar between both uORF sequences.

**Fig. 2:**
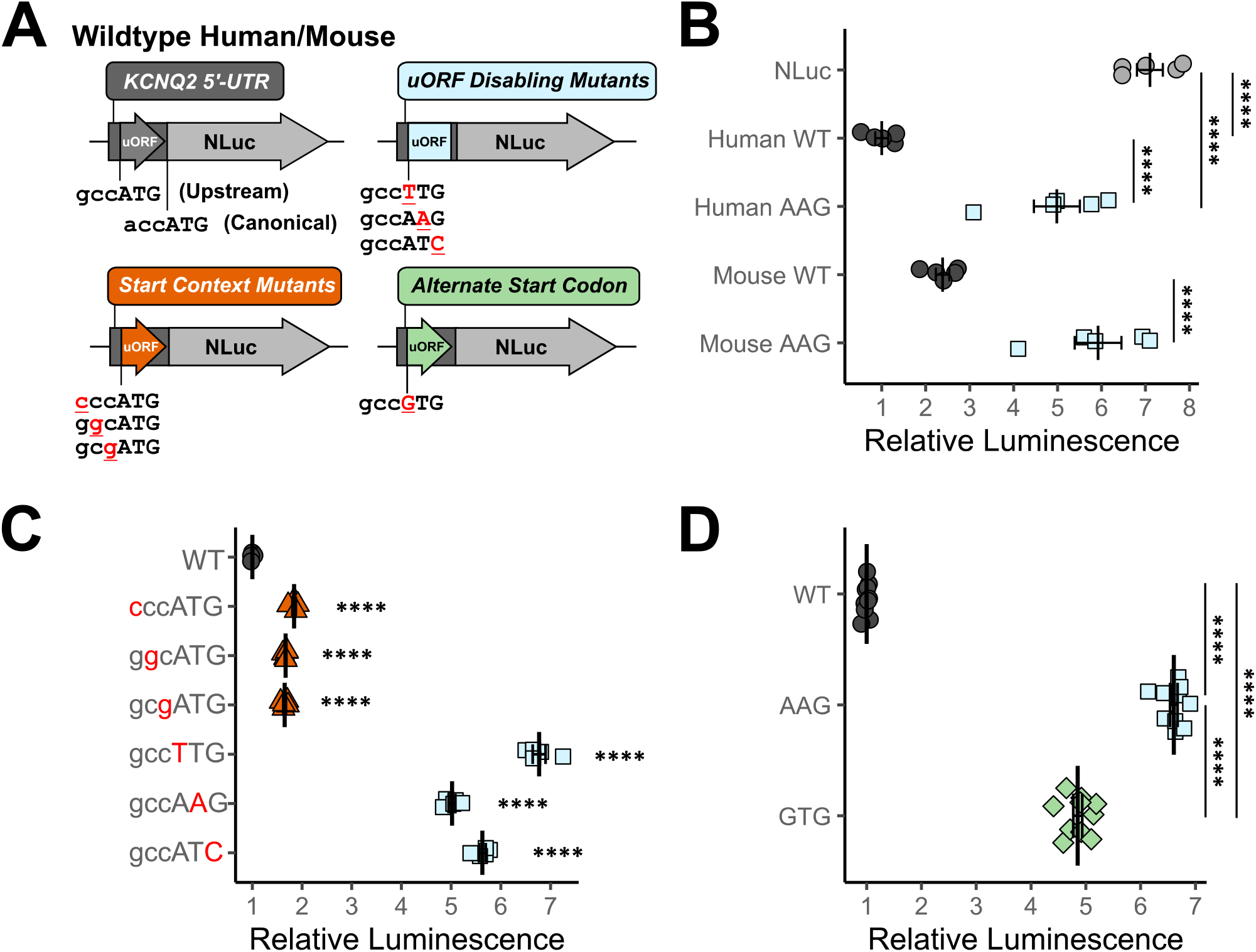
Mutations of the KCNQ2 uORF start codon enhance downstream translation. (**A**) Schematic of *KCNQ2* 5’-UTR-Nluc luciferase constructs. Wildtype (WT) 5’-UTR (dark gray circles) from *KCNQ2* containing the uORF was inserted immediately upstream of the Nluc coding sequence (light gray circles). Site-directed mutagenesis introduced uORF start codon mutations of the complementary base at each position (TTG, AAG, ATC, blue squares). Mutations of the upstream Kozak consensus sequence were also tested (cccATG, ggcATG, gcgATG, orange triangles). A translationally weaker non-cognate start codon was also introduced (GTG, green diamonds). Translational efficiencies with each 5’-UTR mutation are shown for **(B)** human and mouse 5’-UTR and AAG uORF variants compared to the original *Nluc* 5’-UTR (n=5), **(C)** single-nucleotide walk from the upstream start context to the upstream start codon for the human *KCNQ2* 5’-UTR (n=5), and **(D)** GTG and AAG uORF variants (n=9). All experimental data in panels B-D are presented as normalized luminescence (Nluc / Firefly) relative to the WT 5’-UTR condition, measured from HEK293 cells transfected with each construct (Games-Howell post-hoc, **** p<0.00005, mean ± SEM).

To further demonstrate that the upstream ATG truly functions as a start codon, we performed mutational scanning of the region surrounding the ATG codon, focusing on the human *KCNQ2* 5’-UTR. We hypothesized that mutations within the Kozak consensus sequence would weaken the uORF and boost reporter gene translation, with the strongest effect produced through mutation of the purine at the −3 position. Additionally, we predicted that mutations at all three nucleotides of the upstream ATG would enhance translation to a much greater degree. We introduced single nucleotide changes in a 1 nucleotide walk along the uORF including its start codon (ATG) and the three nucleotides upstream (GCC), changing each nucleotide to its complementary base-pair so as not to alter GC content. We observed greater levels of reporter gene translation in all conditions compared to the wildtype sequence (Welch’s ANOVA: p=3.86E-16) (**Fig. 2C**). Mutations at the uORF start codon produced a 5 to 7-fold boost to translation (+1A:T p=5.85E-6, +2T:A p=1.05E-6, +3G:C p=9.83E-7), consistent with previous findings. In addition, mutations upstream of the uORF start codon enhanced translation by only 1.5-fold (−3G:C p=2.44E-5, −2C:G p=2.04E-8, −1C:G p=1.28E-5). Interestingly, the removal of a purine at the −3 position was comparable to mutations at the −1 and −2 position from the upstream ATG, suggesting that the Kozak consensus sequence may not contribute substantially to ribosome initiation at the upstream start site as we hypothesized. We also tested the effect of introducing the translationally weaker near-cognate GTG into the human *KCNQ2* uORF. Compared to the wildtype 5’-UTR, the GTG initiated uORF was associated with higher levels of translation but to a significantly less extent than the AAG mutant uORF (Welch’s ANOVA: p=9.34E-22; WT vs AAG: p=4.73E-11; WT vs GTG: p=1.52E-10, AAG vs GTG p=2.38E-11) (**Fig. 2D**). Notably, no differences were observed with quantification of luciferase RNA between cells expressing WT, AAG, or GTG mutant constructs (Welch’s ANOVA: p=0.26) (**Fig. S1**). Our findings indicate that disrupting the uORF leads to enhancement of canonical translation, consistent with our hypothesis that this 5’-UTR element is a translational repressor.

### Disruption of the *KCNQ2* uORF enhances translation of functional KCNQ2 channels

To understand how the uORF may be involved in the regulation of *KCNQ2*-encoded potassium channels, we assessed uORF-suppression on translation and function of the KCNQ2 protein. We hypothesized that mutation of the uORF would produce more protein and thus greater potassium current density, without altering mRNA levels. We generated plasmids expressing full length *KCNQ2* coding sequence with either wildtype or uORF mutant 5’-UTRs preceding the canonical start codon. To measure functional properties of the potassium channel in its most physiological state, we expressed each plasmid in a CHO cell line stably expressing *KCNQ3* (CHO-Q3) previously generated in our laboratory (39). We measured current density and voltage-dependence of activation using automated planar patch clamp recording. We observed an approximately 2-fold larger peak current density in cells expressing either AAG or GTG uORF mutants, compared to the wildtype 5’-UTR (**Fig. 3A**). At 0 mV, current density was significantly different between conditions (Welch’s ANOVA: p=2.62E-4), with the AAG condition producing 1.9x greater current density (p=0.025) and the GTG condition producing 1.7x greater current density (p=4.89E-4) (**Fig. 3B**). These findings indicate that suppression of the *KCNQ2* uORF within the 5’-UTR enhances current density, consistent with greater numbers of functional channels at the plasma membrane. Voltage-dependence of activation was not affected (Welch’s ANOVA of V_1/2_: p=0.52), consistent with our hypothesis that uORF-suppression produces greater channel expression without altering other biophysical properties (**Fig. 3C**).

**Fig. 3:**
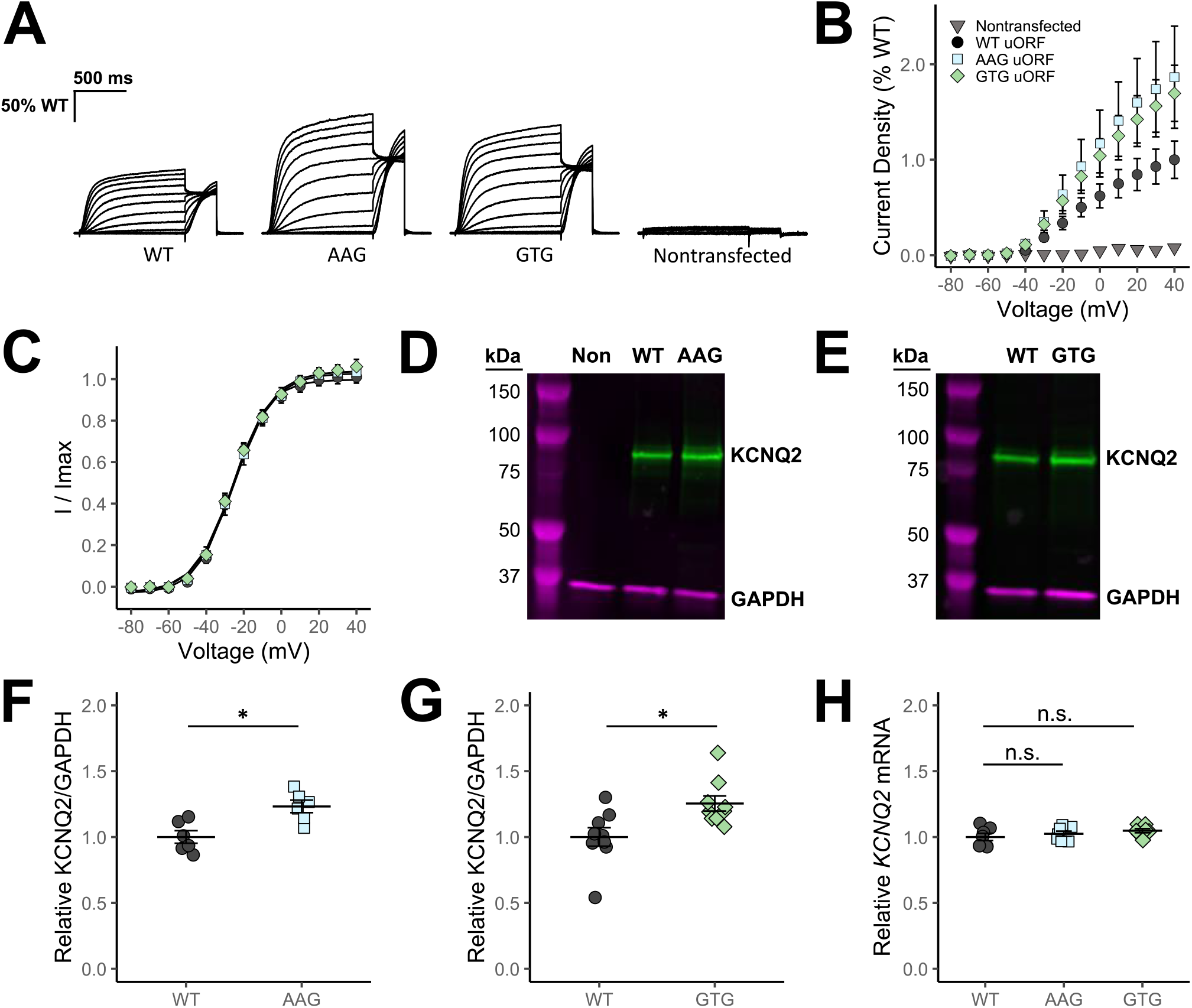
uORF inactivation enhances translation of functional KCNQ2 channels in a post-transcriptional manner. **(A)** Averaged XE-991-sensitive normalized to WT whole-cell current density traces recorded from CHO-Q3 cells electroporated with *KCNQ2* constructs fused to the *KCNQ2* 5’-UTR containing WT, AAG, or GTG uORF mutations. Nontransfected cells are shown as negative control. **(B)** Relationship of average current density and test voltage for each condition (WT: n=105, AAG: n=26, GTG: n=91, Nontransfected: n=41). **(C)** Voltage-dependence of activation measured for each condition (WT: V_1/2_ = −26.1mV, *k* = 9.4, n=54; AAG: V_1/2_ = −25.0mV, *k* = 10.3, n=16; GTG: V_1/2_ = −25.2mV, *k* = 10.5, n=55). **(D)** Representative western blot of KCNQ2 protein (green) and GAPDH (magenta) for CHO-Q3 cell electroporations comparing WT and AAG or **(E)** WT and GTG conditions. **(F)** Quantified KCNQ2 signal normalized to GAPDH for AAG (n=6) and **(G)** for GTG uORF conditions (n=9). **(H)** Relative mRNA levels for *KCNQ2* from CHO-Q3 cells electroporated with each *KCNQ2* construct (Games-Howell post-hoc, n.s. p>0.05, n=9). Scale bars: 500ms (horizontal); 50% WT current (vertical). Data shown in B-C and H are mean ± 95% CI. Data shown in F-G are mean ± SEM (Welch’s t-test, *p<0.05).

To quantify differences in steady-state protein levels, we performed western blot analysis on protein lysates collected from electroporated CHO-Q3 cells. We observed significantly higher KCNQ2 protein levels for cells transfected with AAG and GTG uORF mutant constructs, compared to wildtype cells (**Fig. 3D and 3E**). When normalized to GAPDH as a loading control, 25% greater protein levels were observed for both the AAG mutation (p=6.18E-3) (**Fig. 3F**) and the GTG mutation (p=0.013) (**Fig. 3G**). These results indicate that boosting total KCNQ2 protein levels by 25% correlates with two-fold larger current densities, which supports our hypothesis that uORF-suppression can promote greater expression of KCNQ2 channels.

Suppression of the *KCNQ2* uORF should alter protein translation in a post-transcriptional manner. To determine if the greater protein levels and current density in uORF-inactivated conditions are driven by greater mRNA expression, we quantified steady-state levels of *KCNQ2* mRNA. For both AAG and GTG mutant uORF constructs, *KCNQ2* mRNA levels were not significantly different compared to cells transfected with the wildtype 5’-UTR construct (Welch’s ANOVA: p=0.346) (**Fig. 3H**). We excluded the possibility that greater current density was driven by greater expression of *KCNQ3* after observing no differences in protein (Welch’s ANOVA: p=0.227) (**Fig. S2A and S2B**) or transcript levels for this gene (Welch’s ANOVA: p=0.128) (**Fig. S2C**). We concluded that suppression of the uORF produces greater levels of KCNQ2 channels without altering mRNA expression levels.

### Inactivation of endogenous *KCNQ2* uORF suppresses ribosomal engagement

The previous experiments demonstrate that the *KCNQ2* uORF can be suppressed to promote potassium channel expression in heterologous cells, but it remains unclear whether these findings hold true for the endogenously expressed mRNA. To address this question, we used SH-SY5Y cells, an immortalized human neuroblastoma cell line that endogenously expresses *KCNQ2* at a high level. By contrast, *KCNQ3* is not expressed in SH-SY5Y cells.

To inactivate the *KCNQ2* uORF at the genomic level, we used adenine base editing to convert the upstream ATG start codon to the near-cognate GTG codon. The targeted adenine is an optimal base editing target located in an adenine-poor region, where there are few candidates for bystander edits; the nearest adenines from the targeted nucleotide are located 15 nt 5’ and 65 nt 3’ to the targeted adenine. We introduced a guide RNA sequence capable of directing an adenine base editor (ABE8e) to the uORF start codon into a sgRNA-expression plasmid to be driven by a human U6 promoter. As a negative control, we used a nontargeting guide sequence lacking complementarity with any part of the human genome.

SH-SY5Y cells were electroporated with sgRNA and ABE8e expression plasmids to induce the desired genome edit. The process was repeated three times, producing three replicate edited polyclonal cell lines, each paired with a cell line transfected with the nontargeting sgRNA. Cells transfected with the targeting sgRNA consistently produced high efficiency on-target base editing (Replicate 1: 92% edited, Replicate 2: 91% edited, Replicate 3: 93% edited, mean: 92% edited, standard error = 0.6%, p=2.89E-5 by Welch’s t-test) without bystander edits (**Fig. 4A**). By contrast, the *KCNQ2* uORF in nontargeting controls was not edited.

**Fig. 4:**
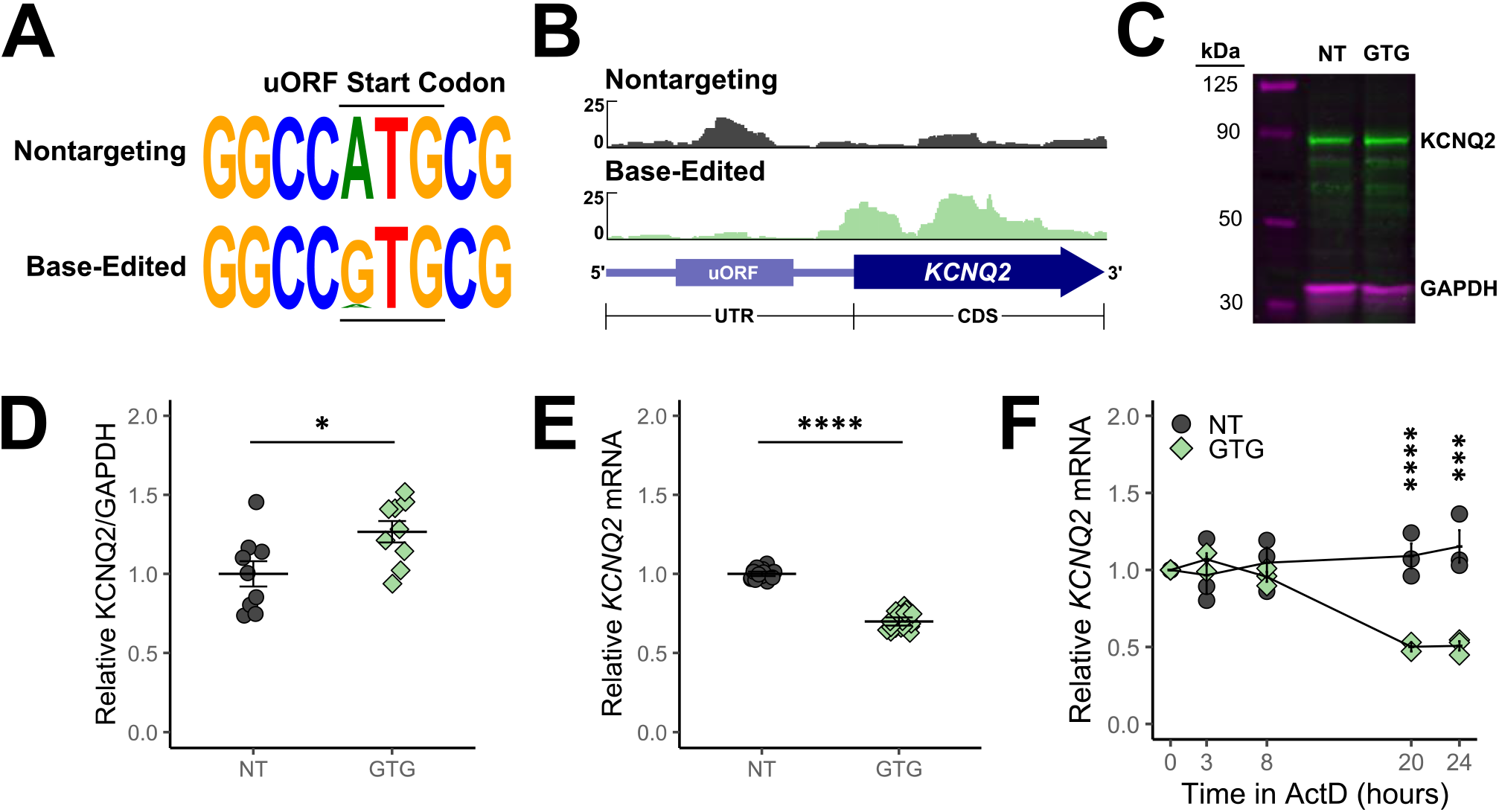
Endogenous inactivation of the *KCNQ2* uORF enhances protein translation, but downregulates *KCNQ2* mRNA levels. **(A)** Sequence logo of the *KCNQ2* uORF start codon region for SH-SY5Y cells electroporated with an adenine base editor and either a nontargeting (NT) or uORF-targeting gRNA. **(B)** Ribosome profiling of the NT (dark gray) and GTG uORF-edited (light green) SH-SY5Y cell lines for the *KCNQ2* 5’-UTR and 5’-proximal CDS region. Ribosome footprints are shown as combined signal from three independently electroporated cell lines per condition. **(C)** Representative western blot for KCNQ2 (green) and GAPDH (magenta) for NT and GTG uORF-edited SH-SY5Y cells. **(D)** Quantification of KCNQ2 signal normalized to GAPDH, relative to the NT condition of each electroporation (Welch’s t-test, * p<0.05, mean ± SEM, n=3 per replicate cell line, n=9 per condition).Relative *KCNQ2* mRNA levels for NT and GTG uORF-edited SH-SY5Y cells, relative to the NT condition of each electroporation (Welch’s t-test, **** p<0.00005, mean ± 95% CI, n=4 per replicate cell line, n=12 per condition). **(F)** Time-course of relative *KCNQ2* mRNA decay for NT and GTG uORF-edited SH-SY5Y cells following treatment in actinomycin D, normalized to the 0-hour time point of each condition. (Welch’s t-test, *** p<0.0001, **** p<0.00001, mean ± SEM, n=3 independent experiments)

We hypothesized that in the edited cell lines, ribosome occupancy at the *KCNQ2* uORF would be lower compared to non-edited cells. To investigate this, we treated each cell line with retinoic acid to promote neuronal differentiation, then performed ribosome profiling using RNase footprinting. Analysis of ribosome footprints at the *KCNQ2* uORF revealed that edited cells exhibited lower ribosome occupancy at the uORF compared to the nontargeting controls (**Fig. 4B**). Edited cells also showed greater ribosome occupancy at the canonical *KCNQ2* ORF consistent with a greater level of translation. As a negative control, no difference in uORF translation was observed between conditions for the single uORF gene *RNASEH1* (40), suggesting that the effect is specific to *KCNQ2* (**Fig. S3**).

### Inactivation of endogenous KCNQ2 uORF enhances protein expression but downregulates steady-state mRNA levels

Given our findings from *KCNQ2* overexpression experiments, we hypothesized that uORF-edited cells would produce greater protein expression compared to non-edited cells, with no change in mRNA transcript expression. By western blot analysis, we observed 30% greater KCNQ2 protein levels in edited cells (**Fig. 4C and D**). This was consistent with findings from transient transfection, confirming that the effect holds true when inactivating the endogenous uORF. However unexpectedly, steady-state levels of *KCNQ2* mRNA were 30% lower in edited cells (**Fig. 4E**).

One explanation for lower steady-state RNA levels is that the mRNA stability is altered, leading to faster mRNA decay. To assess whether uORF-suppression alters *KCNQ2* mRNA decay, we treated one edited cell line and its nontargeting control with actinomycin D (ActD) to disable transcription, then measured *KCNQ2* mRNA over time. Across three independent experiments, we observed that *KCNQ2* mRNAs from nonedited cells do not undergo measurable decay within 24 hours of ActD treatment, but in uORF-edited cells, mRNA levels fall to half of the starting amount by 20 hours (p=6.21E-5) (**Fig. 4F**). We concluded that the GTG mutation of the *KCNQ2* uORF destabilizes the mRNA transcript, resulting in lower steady-state mRNA compared to nonedited cells.

## Discussion

Regulatory regions in the 5’-UTR of clinically relevant genes are often understudied. Here, we show for a potassium channel gene associated with monogenic epilepsy (*KCNQ2*), that a single uORF present within the 5’-UTR represses translation of the encoded protein. Additionally, disabling the uORF start codon boosts translation of potassium channels. These findings demonstrate a regulatory feature within the 5’-UTR of a disease-associated gene, revealing insights for control of *KCNQ2* translation and potentially for therapeutic targeting.

Although the *KCNQ2* uORF can be inactivated to enhance translation of the protein, the physiological function of the *KCNQ2* uORF remains unclear. It is known that for some genes that the peptide product from uORF translation is stable and functional (41–44). However, uORF peptides shorter than 30 amino acids are generally unstable (45), and functional predictions based on existing proteomics data offer weak evidence for a stable peptide encoded by the *KCNQ2* uORF (4). Given the length and low prediction score for the *KCNQ2* uORF peptide product, we believe that this uORF product is not likely to be stable.

Aside from producing peptides, uORFs are known to act as translational switches under different conditions (9, 46, 47). The most well-studied example is the *GCN4* gene, where cellular stress induces translation of the protein despite global repression of translation (48, 49). However, this stress-responsive activity occurs by a multi-uORF system through a delayed-reinitiation mechanism (50), which would not apply to single-uORF genes (15). Given that *KCNQ2* possesses a single uORF, we posit that it is less likely to be regulated by stress.

For neuronal genes, emerging evidence suggests that uORFs may act to fine-tune translation in an activity-dependent manner. Translation is required for synaptic plasticity during which localized protein translation at synapses undergo rapid remodeling following cellular excitation (51, 52). The relation to uORF translation is less well understood but has recently emerged as a point of interest in the field. In one study evaluating the spatial regulation of activity-dependent translation in primary cortical neurons, depolarization promoted uORF translation selectively in dendrites (53). In another study modeling short and long-term effects of activity-dependent translation in SHSY5Y cells, mRNA dynamics of genes containing upstream ATG codons were altered immediately after depolarization, but resynchronized after a recovery period of just two hours (54). We interrogated these published data to determine whether translation of the *KCNQ2* uORF is altered after depolarization. We observed greater uORF translation for *KCNQ2* immediately after depolarization compared to cells at rest, but this effect was absent following the two-hour recovery period (**Fig. S4**). In contrast, total mRNA levels for *KCNQ2* were not changed. These observations may implicate the *KCNQ2* uORF as a regulatory feature during activity-dependent translation, but the exact physiological purpose of this feature remains unknown.

Interestingly, here we report that inactivation of the *KCNQ2* uORF in differentiated SH-SY5Y cells results in a faster mRNA decay rate and lower steady-state mRNA levels. This is counter-intuitive to what has been previously reported for uORF-containing genes, in which uORF stop codons are commonly recognized as premature terminations, triggering the mRNA surveillance pathway known as nonsense-mediated decay (NMD) (55, 56). It is known that premature termination codons close to an exon-exon junction are able to escape NMD (57), but this is located beyond the reported 55 nt threshold for *KCNQ2*, suggesting that the native transcript should be a target for NMD. It is possible that the *KCNQ2* uORF is too short to engage NMD, which is similar to the NMD-insensitive gene *THPO* (58). This could explain why *KCNQ2* mRNA levels are not higher after mutation of the uORF start codon, but it does not explain why they are lower.

Faster *KCNQ2* mRNA decay when its uORF is disabled could be a form of compensation. For example, to prevent KCNQ2 protein overproduction from transcripts lacking the uORF, a separate mRNA decay pathway may be recruited to downregulate *KCNQ2* at the mRNA level. Alternatively, suppression of the uORF could directly destabilize the mRNA. It is conceivable that the uORF serves as a translational buffer (59), and that transcripts lacking the uORF are predisposed to more ribosome collisions, triggering the no-go decay (NGD) pathway to degrade the mRNA (60). It is unclear why *KCNQ2* mRNA was not lower in transient transfection systems, but perhaps its decay is maximized under overexpression, masking physiological decay rates. Limiting the system to a single transcript isoform of *KCNQ2* could also obscure detection of physiological mRNA dynamics. Regardless, these experiments demonstrate the importance of assessing uORF function through physiological suppression at the genetic level.

Targeting the *KCNQ2* uORF could serve as a potential therapeutic strategy for upregulating KCNQ2 protein in the setting of heterozygous loss-of-function variants. In our study, we demonstrated that adenine base editing of the upstream ATG start codon can suppress uORF translation and boost translation of the canonical reading frame, similar to other studies (16). The *KCNQ2* uORF start codon also represents an ideal base editing target, due to the lack of bystander adenosine nucleotides. Theoretically, this single nucleotide edit would provide a therapeutic strategy for loss-of-function *KCNQ2* variants associated with haploinsufficiency.

### Study Limitations

Although the effects of this uORF are relatively consistent across the model systems used in our study, there are experimental limitations. Heterologous expression systems may not accurately represent native cell types. For example, the stoichiometry of KCNQ2 and KCNQ3 subunits may be skewed towards *KCNQ2*, which is transiently expressed, relative to the single copy of stably expressed *KCNQ3*. However, we previously demonstrated absence of KCNQ2 homotetrameric channel assemblies based on the pharmacological sensitivity of recorded ion currents (39). A similar concern exists for SH-SY5Y cells, which do not express *KCNQ3* at measurable levels. However, these cells are a suitable model for assessing base editing given their near diploid karyotype, neuronal lineage, and ease of transfection. By comparison, delivery of base-editing machinery to iPSC-derived neurons is challenging and inefficient (61).

The therapeutic value of disabling the *KCNQ2* uORF may be limited in some situations. Notably, KCNQ2 protein expression in both heterologous cells and base edited cells rose by 30%, which is a substantially lower amount than the 6-fold boost observed in reporter gene assays. However, this level of enhanced protein expression was still physiologically impactful given that current densities nearly doubled, which may reflect a disproportional share of channel protein reaching the plasma membrane where ion conductance occurs. Additionally, the relative level of ribosome engagement observed at the human *KCNQ2* uORF appears lower than at the canonical translation start site (**Fig. 1B**), which suggests a potential limitation of the therapeutic impact of suppressing this element. However, uORF translation efficiency may vary by brain region. Further studies to determine the relative level of uORF versus canonical translation in different brain regions may help anticipate the therapeutic potential of targeting the *KCNQ2* uORF.

Importantly, a uORF suppression strategy targets transcripts from both alleles, including the disease-associated allele. While this may be effective for variants causing haploinsufficiency, it may be less impactful for *KCNQ2* variants that exhibit dominant-negative effects. Loss-of-function variants causing haploinsufficiency are more frequent in SLFNE, although an estimated 10-15% of these individuals develop seizures later in life suggesting they could benefit from KCNQ2 upregulation (62). By contrast, dominant-negative variants are more frequently associated with DEE, and targeting the *KCNQ2* uORF in this setting is predicted to be less effective (63). There may be an opportunity for coupling uORF suppression with allele-specific knockdown of the variant transcript to restore normal levels of KCNQ2 protein in all situations. A promising approach to accomplish this relies on siRNA or gapmer ASO technology to specifically degrade RNA from the variant allele (64, 65). However, this requires an individualized approach. Our strategy to upregulate KCNQ2 expression by targeting the uORF would be most meaningful for variants that are not dominant-negative.

In summary, we report that the *KCNQ2* 5’-UTR contains a uORF sequence that represses KCNQ2 protein translation. Our work highlights an element within the 5’-UTR that can be otherwise overlooked in disease-associated genes.

## Acknowledgements

We thank Daniel Arango for insightful comments and reading of this manuscript. We thank Christopher Thompson, Jennifer Kearney, and Evangelos Kiskinis for helpful discussions, as well as Naiyu Shi for technical assistance with ribosome profiling. This work was supported in part by NIH grants NS137587 and OD034362. D.J.H. is supported by a T32 Predoctoral Training Grant (T32GM105538) from the National Institute of General Medical Sciences and a National Research Service Award (NRSA) fellowship from the National Institute of Neurodevelopmental Disorders and Stroke (F31NS135753). E.G. is supported by a postdoctoral fellowship from the American Heart Association (26POST1542927). Z.J. was supported by NIH grants R01HL161389 and R35GM138192.

## Author contributions

D.J.H., Z.J., A.L.G. designed research; D.J.H., E.G., C.G.V., C.Q.S., Q.L. performed research; J.-M.D.K. contributed new reagents; D.J.H., E.G., J.-M.D.K., C.G.V., E.K.S. analyzed data; D.J.H., A.L.G. wrote the paper.

## Materials and Methods

### Comparative genomics

Orthologous nucleotide sequences for the *KCNQ2* 5’-UTR were collected from the NCBI nucleotide database for the following species: human (Homo sapiens, NM_004518.6), macaque (Macaca mulatta, XM_015148615.2), mouse (Mus musculus, NM_010611.3), rat (Rattus norvegicus, NM_133322.2), whale (Delphinapterus leucas, XM_022597496.1), dog (Canis lupus familiaris, NC_051828.1), hamster (Cricetulus griseus, XM_027419933.1), turtle (Chrysemys picta bellii, XM_005309569.4), lungfish (Protopterus annectens, XM_044090449.1), and zebrafish (Danio rerio XM_009302767.5). Multiple sequence alignment was performed using the Clustal Omega web server (66).

### Ribosome Profiling

For human and mouse brain Ribo-seq datasets, reads were collected from Sequencing Reads Archive (SRA) Accession PRJNA223060 (36). Ribo-seq datasets for depolarized SHSY5Y cells were collected from SRA Accession PRJNA655417 (54). For obtaining Ribo-seq datasets from base edited SHSY5Y cells, sequencing libraries were prepared according to Li et al (67). Briefly, cells were pre-treated with cycloheximide (100µg/mL) for 5 minutes at 37°C to halt actively translating 80S translation complexes. Cell lysates were then prepared and treated with 0.526U/µL RNase I (LGC Biosearch Technologies) for 1.5 hours with gentle agitation at room temperature. RNase footprints were recovered using TRIzol (Invitrogen), followed by end-healing with T4 polynucleotide kinase (New England Biolabs) and polyadenylation by *E. coli* poly(A) polymerase (New England Biolabs). The SMART-RT approach (68) was used to generate cDNA libraries with partial Illumina sequencing adapters through Superscript II reverse transcriptase (New England Biolabs) primed with oligo(dT) primer and template switching using a SMARTer oligo (**SI Appendix, Table S1**). Complete adapter sequences and indices (**SI Appendix, Table S2**) were added in two rounds of PCR using Q5 DNA polymerase (New England Biolabs). Libraries were pooled and sequenced using an Illumina Novaseq X Plus platform (Admera Health) to obtain at least 50 million paired-end clusters per sample. For data analysis, reads were trimmed according to the poly(A) sequence or reported adapter sequence with CutAdapt (v4.2), then rRNA-mapped reads were removed using Bowtie2 (v2.4.1). Remaining reads were mapped to the corresponding human or mouse transcriptome using TopHat (v2.1.0). Sequence coverage tracks of uniquely mapped reads were produced for each sample using deepTools (v3.1.1) and visualized in the Integrative Genome Viewer (v2.17.4).

### Plasmids and mutagenesis

Human and mouse *KCNQ2* 5’-UTR were amplified from HEK293 and C57BL/6J genomic DNA, respectively, using PCR with Q5 DNA polymerase (New England Biolabs). Amplicons were cloned into the 5’-UTR position of the pNL1.1.CMV[Nluc/CMV] plasmid (Promega) using NEBuilder HiFi DNA Assembly (Promega). For functional studies, the wildtype *KCNQ2* 5’-UTR sequence was cloned into the 5’-UTR position of the sCMV-KCNQ2-IRES-EGFP plasmid vector generated previously (69). Plasmids were used to transform chemically competent Top10 cells, which were then cultured in Luria-Bertani broth (Invitrogen) containing 100µg/µL ampicillin (Sigma). Plasmids were prepared using the NucleoSpin Plasmid kit (Macherey Nagel). Site-directed mutagenesis was performed using Q5 PCR as previously described (70). All cultures were grown for large-scale plasmid preparation and verified using nanopore whole-plasmid sequencing (Plasmidsaurus, Inc). Endotoxin-free DNA was prepared using the NucleoBond Xtra Maxi EF kit (Macherey Nagel). All primer sequences are provided in the SI Appendix, Table S2.

### Cell culture

All cell lines were obtained initially from the American Type Culture Collection and cultured at 37°C in 5% CO_2_ in varying base media. HEK293 cells (CRL-1573) were cultured with Dulbecco’s Modified Eagle Medium (Gibco), CHO-K1 cells (CCL-61) were cultured with Kaighan’s F12 medium (Gibco), and SH-SY5Y cells (CRL-2266) were cultured with DMEM/F12 medium containing GlutaMAX (Gibco). For functional studies, CHO-K1 cells stably expressing *KCNQ3* (CHO-Q3) previously generated in our laboratory (39) were cultured in Ham’s F12 media (Gibco) and 600 μg/mL hygromycin B (Invitrogen) to maintain selection. All media were supplemented with 10% fetal bovine serum (Atlanta Biologicals), 50 units/mL penicillin (Gibco), 50 μg/mL streptomycin (Gibco), and 2 mM L-glutamine (Gibco) if not already containing GlutaMAX.

### Reporter gene assays

Cells were plated at a density of 10,000 cells per well of a 24-well plate, 24 hours prior to transfection. Cells were co-transfected with NanoLuc-expressing constructs and pGL4.50[*luc2*/CMV/Hygro] (Promega) encoding Firefly luciferase using FuGENE 6 (Promega) following manufacturer’s guidelines. Twenty-four hours after transfection, cells were lysed in passive lysis buffer (Promega). Lysates were diluted 1:50 in water and transferred to wells of a white 96-well plate (Greiner Bio-One). Luminescence from both enzymes were induced using the Nano-Glo Dual Luciferase Reporter Assay System (Promega) following manufacturer’s guidelines and measured using a Varioskan LUX plate reader (Thermo Fisher Scientific). Relative luminescence was determined by normalizing the signal from the NanoLuc luciferase with that from the Firefly luciferase.

### Electroporation of KCNQ2-expressing plasmids

Cells were electroporated according to the previously published methods (71, 72). CHO-Q3 cells were cultured to 70-80% confluence before trypsinization with TrypLE (Thermo Fisher Scientific). Viable cell concentration of the cell suspension was determined using an automated cell counter (ViCell, Beckman Coulter). Cell pellets were obtained by centrifugation at 193 × g for 4 min and gently washed with 5 mL of electroporation buffer (EBR100; MaxCyte, Inc.) before centrifuging again (193 × g, 2 mins). The final cell pellets were resuspended in a small volume of electroporation buffer to a final density of 10^8^ viable cells/mL.

Plasmid DNA (20 µg) containing the native or mutant *KCNQ2* 5’-UTRs was mixed with 100 μL of cell suspension (10^7^ viable cells). The DNA-cell suspension was transferred to an OC-100×2 processing assembly (MaxCyte, Inc.) and electroporated using the MaxCyte STX system (MaxCyte, Inc.) with the “CHO2 (Protein Expression)” protocol. Immediately after electroporation, 10 μL of recombinant human DNaseI (Pulmozyme, Genentech, Inc.) was added to the electroporated cells, followed by recovery in a 60 mm dish at 37°C and 5% CO_2_ for 30 minutes. After recovery, cells were resuspended in complete media without hygromycin and cultured in a 10 cm dish for 24 hours. Cells were harvested and frozen in complete media supplemented with 10% DMSO (Sigma), in 1 mL aliquots of 1.8×10^6^ viable cells/mL. Cells were thawed 30 hours prior to western blot, quantitative PCR, and electrophysiology experiments, and incubated at 37°C and 5% CO_2_. Twenty hours prior to experiments, cells were incubated at 28°C and 5% CO_2_ to promote KCNQ2 channel migration to the cell surface.

### Western blot

Cells were collected in ice-cold PBS (Thermo Fisher Scientific) containing Halt protease and phosphatase inhibitor cocktail (Thermo Fisher Scientific) using a cell scraper, then pelleted at 5687 × g for 4 minutes. Cell pellets were lysed in RIPA buffer (Thermo Fisher Scientific) supplemented with the protease and phosphatase inhibitor cocktail for 1 hour at 4°C with gentle agitation, then centrifuged at 12000 × g for 10 minutes. Supernatants were collected for western blotting. Protein concentrations were determined using the Pierce BCA Assay (Thermo Fisher Scientific) microplate protocol. Protein samples (5 μg for CHO-Q3, 30 μg for SH-SY5Y) were combined with loading buffer (LiCor) and 50 mM dithiothreitol (Bio-Rad), then incubated at room temperature for 30 minutes. Samples were separated by SDS PAGE using the Mini-PROTEAN electrophoresis system (Bio-Rad). Samples loaded on a 7.5% Mini-PROTEAN TGX pre-cast polyacrylamide gel (Bio-Rad) were electrophoresed for 1 hour at 100V in Tris-glycine running buffer (25 mM Tris, 192 mM glycine, 0.1% SDS). Proteins were transferred to Immobilon PVDF membrane (Millipore) re-hydrated with 100% methanol (Sigma) using the Mini-PROTEAN mini trans-blot module. Transfer was performed overnight at 4°C with constant 15V from an Owl EC-105 power supply (Thermo Fisher Scientific) in a Tris-glycine transfer buffer (25 mM Tris, 192 mM glycine, 20% methanol).

Membranes were washed three times in TBS-T (20 mM Tris, 150 mM NaCl, 0.1% Tween-20) before blocking for 1 hour with 5% blotting-grade milk (Bio-Rad) in TBS-T with gentle agitation. Primary antibody incubation was performed overnight at 4°C with gentle agitation using rabbit anti-KCNQ2 (14752S, Cell Signaling Technologies, 1:1000 dilution), or rabbit anti-KCNQ3 (APC-051, Alomone labs, 1:1000 dilution), and mouse IgG1a anti-GAPDH (AM4300, Invitrogen, 1:7500 dilution). Membranes were washed three times in TBS-T, then incubated for 1 hour with gentle agitation using 800CW goat anti-rabbit (LiCor, 1:20000 dilution) and 680LT goat anti-mouse IgG1a (LiCor, 1:50000 dilution) fluorescent secondary antibodies. Membranes were again washed three times in TBS-T, then imaged using an Odyssey CLx scanner (LiCor). Band intensities were quantified using ImageJ (73). All buffer components were obtained from Sigma-Aldrich, except for Tris and Tween-20, which were obtained from Fisher Scientific.

### Quantitative RT-PCR

Total RNA was extracted from cells using TRIzol (Invitrogen), and cDNA was synthesized using SuperScript IV reverse transcriptase (Thermo Fisher Scientific) with random hexamers. For time-course studies to determine mRNA decay rate, cells were first treated with 10 μg/mL Actinomycin D (Sigma Aldrich) for 0, 3, 8, 20, and 24 hours, followed by RNA extraction. Quantitative PCR was performed using the PrimeTime master mix (Integrated DNA Technologies). For expression experiments in CHO-Q3 cells, PrimeTime probe-based qPCR assays (Integrated DNA Technologies) were obtained for human *KCNQ2* (Hs.PT.58.39257585, 6-FAM fluorophore), human *KCNQ3* (Hs.PT.58.3591262, Cy5 fluorophore), and custom assays were designed for endogenous hamster *B2M* as well as luciferase targets (**SI Appendix, Table S3**). For experiments in SH-SY5Y cells, a human *B2M* assay (Hs.PT.58v.18759587, HEX fluorophore) was used as an internal control. All probe assays contained ZEN / Iowa Black FQ as an internal and 3’ quencher. Reactions were performed using a QuantStudio 7 Flex thermocycler (Thermo Fisher Scientific) using default thermal cycling parameters (1 cycle: 50°C, 2 min; 95°C 10 min. 40 cycles: 95°C 15 sec; 60°C 1 min). Relative mRNA levels were calculated using the ΔΔCt method, normalized to *B2M*, or to *Fluc* for luciferase experiments.

### Automated patch clamp recording

Automated patch clamp recordings were performed on electroporated CHO-Q3 cells using the Syncropatch 384 (Nanion Technologies) using the procedure adapted from Vanoye et al (71). Recordings were performed using single-hole 384-well S-type patch plates (Nanion Technologies). The composition of the external solution was 140 mM NaCl, 4 mM KCl, 2 mM CaCl_2_, 1 mM MgCl_2_, 10 mM HEPES, and 5 mM glucose, with pH adjusted to 7.4 using NaOH and osmolarity adjusted to 298 mOsm/kg. The composition of the internal solution was 60 mM KF, 60 mM KCl, 10 mM NaCl, 10 mM HEPES, 10 mM EGTA, and 5 mM MgATP, with pH adjusted to 7.2 using KOH and osmolarity adjusted to 285 mOsm/kg. Whole-cell currents were recorded from −80mV to +40mV in 10mV steps. Nonspecific currents were recorded following treatment with the M-channel blocker XE-991 (Tocris) at 30 µM and digitally subtracted from the total current recorded at each voltage step to determine channel-specific current. All chemicals, unless otherwise indicated, were obtained from Sigma-Aldrich.

Data from each recorded cell were filtered according to stringent cutoff thresholds (before blocker: seal resistance ≥ 250 MΩ, series resistance ≤ 25 MΩ, capacitance ≥ 2 pF, minimum baseline current < −400 pA; after blocker: seal resistance ≥ 200 MΩ, series resistance ≤ 35 MΩ, capacitance ≥ 2 pF, minimum baseline current < −400 pA). Additionally, data were excluded from analysis if there was a run-up of peak current amplitude between treatments that exceeded 15%. Cells with less than 10% of the mean peak current density of each condition were defined as not transfected and excluded from analysis. For voltage-dependence of activation, samples were excluded from analysis if the goodness of fit for a Boltzmann fit was less than 0.8 (R^2^ < 0.8).

### Adenine base editing

A vector for expressing sgRNA (pSpCas9(BB)-2A-GFP (PX458)) was obtained as a generous gift from Feng Zhang (Addgene plasmid #48138). The vector was re-engineered to remove the Cas9 sequence while retaining EGFP. Specifically, PCR using Q5 DNA polymerase (New England Biolabs) was used to amplify the EGFP sequence, followed by digestion of the amplicon and backbone with AgeI and EcoRI (New England Biolabs). The digested fragments were then ligated using T4 DNA ligase (Promega), creating plasmid psgRNA-EGFP. The plasmid encoding the ABE8e adenine base editor fused to an spCas9 nickase was obtained as a generous gift from David Liu (Addgene Plasmid #138489).

For sgRNA design, guide sequences targeting the *KCNQ2* uORF were screened from a list of NGG protospacer motifs at the locus reported from the CRISPOR web server, such that the adenosine targeted for editing was within 12 to 18 nucleotides upstream of the PAM site. The guide sequence chosen for base editing (GCCATGCGGCTCCCGGCCGG) included the target adenine 17 nucleotides upstream of the PAM site with no other adenines in the theoretical base editing window. As a negative control, a non-targeting guide sequence (CGTTAATCGCGTATAATACGG) was obtained from the Cas12a negative control sequence #1 reported by Integrated DNA Technologies, which was further validated in the CRISPOR web server to show no predicted off-target sites in the human genome for a Cas9 NGG PAM motif. Single-stranded DNA oligomers encoding each target sequence in both sense and antisense directions were synthesized (Thermo Fisher Scientific) with the 5’ end appended with sequences corresponding to overhangs generated from BbsI digestion (New England Biolabs). For the non-targeting guide sequence, an additional guanosine (sense) / cytidine (antisense) base pair was added immediately upstream to ensure maximal transcriptional efficiency with the U6 promoter. Oligonucleotides corresponding to each guide were then hybridized at 95°C for 15 mins in nuclease-free duplex buffer (Integrated DNA Technologies), then the resulting inserts were cloned into 25 ng of BbsI-digested psgRNA-EGFP plasmid vector, followed by ligation using T4 DNA ligase. Plasmids were then cloned following methods described in the Plasmids and Mutagenesis section.

SH-SY5Y cells were grown to 90% confluence on the day of transfection. ABE8e plasmid (30 µg) and U6-sgRNA encoding plasmids (10 µg) were electroporated into SH-SY5Y cells using the MaxCyte STX electroporation system with Optimization 3. Cells were grown in culture for 7 days before harvesting and frozen at 3×10^6^ viable cells/mL. For experiments, SH-SY5Y cells were cultured for 5-8 days in 10 mM all-trans retinoic acid (Thermo Scientific Chemicals) to promote neuronal differentiation.

To evaluate editing efficiency, 0.5×10^6^ cells were pelleted at 193 × g for 4 mins, then lysed with lysis buffer (10 mM Tris, pH 7.4, 1% SDS, 10 mM EDTA, 100 mM NaCl) containing 260 μg/mL proteinase K (Sigma Aldrich) overnight at 50°C. Genomic DNA was extracted using a phenol-chloroform method. A phenol chloroform isoamyl alcohol mixture (Sigma Aldrich) was added to lysates, then centrifuged at 20627 × g for 10 minutes to separate layers. The aqueous layer was extracted, and DNA was precipitated with twice the volume of 100% ethanol (Sigma Aldrich) before pelleting at 20627 × g for 10 minutes. Pellets were washed gently with 70% ethanol, decanted, and air-dried. Pellets were resuspended in endotoxin free TE-buffer (10 mM Tris/ HCl, pH 7.5, 1 mM EDTA, Macherey Nagel). The *KCNQ2* uORF region was amplified from genomic DNA using Q5 PCR. Amplicons were electrophoresed in a 2.5% agarose/TAE gel, then gel purified using the Gel and PCR Purification Kit (Macherey Nagel) following manufacturer’s directions. Purified amplicons were sent for Premium PCR nanopore sequencing (Plasmidsaurus, Inc.) and editing efficiency was calculated from the raw reads using the CRISPResso2 web server (74). A sequence logo for showing editing efficiency was created using the WebLogo3 web server (75), following local alignment and trimming of the reads at the editing window.

### Statistical analysis

All data analysis was performed using the R statistical analysis software (76), with plots generated using the ggplot2 package (77). Pairwise comparisons of means were performed using Welch’s t-tests. For comparisons of more than two groups, Welch’s ANOVA was performed with post-hoc analysis by the Games-Howell method. Data points exceeding two times the standard deviation from the mean in any condition were excluded as outliers. All comparisons were performed using a significance threshold of α=0.05, with Bonferroni correction applied for multiple tests.

## References

1. B. M. Pickering, A. E. Willis, The implications of structured 5’ untranslated regions on translation and disease. Semin. Cell Dev. Biol. 16, 39–47 (2005).

2. J. D. French, S. L. Edwards, The role of noncoding variants in heritable disease. Trends Genet. 36, 880–891 (2020).

3. A. V. Kochetov, A. Sarai, I. B. Rogozin, V. K. Shumny, N. A. Kolchanov, The role of alternative translation start sites in the generation of human protein diversity. Mol. Genet. Genomics 273, 491–6 (2005).

4. P. McGillivray, et al., A comprehensive catalog of predicted functional upstream open reading frames in humans. Nucleic Acids Res. 46, 3326–3338 (2018).

5. K. Wethmar, The regulatory potential of upstream open reading frames in eukaryotic gene expression. Wiley Interdiscip. Rev. RNA 5, 765–78 (2014).

6. C. Barbosa, I. Peixeiro, L. Romão, Gene expression regulation by upstream open reading frames and human disease. PLoS Genet. 9, e1003529 (2013).

7. Y. Kurihara, Uorf shuffling fine-tunes gene expression at a deep level of the process. Plants 9, 1–7 (2020).

8. X. Jin, E. Turcott, S. Englehardt, G. J. Mize, D. R. Morris, The two upstream open reading frames of oncogene mdm2 have different translational regulatory properties. J. Biol. Chem. 278, 25716–21 (2003).

9. N. Ghilardi, A. Wiestner, R. C. Skoda, Thrombopoietin production is inhibited by a translational mechanism. Blood 92, 4023–30 (1998).

10. G. Loughran, et al., Unusually efficient CUG initiation of an overlapping reading frame in POLG mRNA yields novel protein POLGARF. Proc. Natl. Acad. Sci. U. S. A. 117, 24936–24946 (2020).

11. X.-H. Liang, et al., Translation efficiency of mRNAs is increased by antisense oligonucleotides targeting upstream open reading frames. Nat. Biotechnol. 34, 875–80 (2016).

12. L. Zhang, et al., Antisense oligonucleotide-mediated upregulation of Jag1 ameliorates liver disease phenotypes in a mouse model of Alagille syndrome. Mol. Ther. Nucleic Acids 36, 102694 (2025).

13. N. Ahlskog, et al., uORF-targeting steric block antisense oligonucleotides do not reproducibly increase RNASEH1 expression. Mol. Ther. Nucleic Acids 36, 102406 (2025).

14. M. Doisy, et al., Targeting a pathogenic variant creating an upstream AUG in the ENG 5’ untranslated region with antisense oligonucleotides fails to restore protein expression. Nucleic Acid Ther. 36, 12–22 (2026).

15. L. Chen, X. Gao, X. Liu, Y. Zhu, D. Wang, Translational regulation of PKD1 by evolutionarily conserved upstream open reading frames. RNA Biol. 22, 1–12 (2025).

16. N. Feng, et al., Targeted BDNF upregulation via upstream open reading frame disruption. Mol. Ther. 34, 1652–1671 (2026).

17. H. Yan, W. Tang, Programmed RNA editing with an evolved bacterial adenosine deaminase. Nat. Chem. Biol. 20, 1361–1370 (2024).

18. O. M. Hedaya, et al., Secondary structures that regulate mRNA translation provide insights for ASO-mediated modulation of cardiac hypertrophy. Nat. Commun. 14, 6166 (2023).

19. Z. Ang, et al., Alternative splicing of its 5’-UTR limits CD20 mRNA translation and enables resistance to CD20-directed immunotherapies. Blood 142, 1724–1739 (2023).

20. S. de Breyne, et al., Alternative splicing in the RBMXL1 5’-UTR induces uORF-mediated translation control in activated B lymphocytes. Sci. Rep. 15, 22800 (2025).

21. V. L. Romanelli Tavares, et al., Craniofrontonasal Syndrome Caused by Introduction of a Novel uATG in the 5’UTR of EFNB1. Mol. Syndromol. 10, 40–47 (2019).

22. S. R. F. Twigg, et al., Cellular interference in craniofrontonasal syndrome: males mosaic for mutations in the X-linked EFNB1 gene are more severely affected than true hemizygotes. Hum. Mol. Genet. 22, 1654–62 (2013).

23. T. Hayashi, et al., Identification of 5’ untranslated region variants in genes involved in neurodevelopmental disorders. J. Hum. Genet. 1–9 (2026). 10.1038/s10038-025-01446-7.

24. S. Weckhuysen, A. L. George Jr., KCNQ2- and KCNQ3-associated epilepsy (Cambridge University Press, 2022).

25. N. M. Allen, S. Weckhuysen, K. Gorman, M. D. King, H. Lerche, Genetic potassium channel-associated epilepsies: clinical review of the Kv family. Eur. J. Paediatr. Neurol. 24, 105–116 (2020).

26. C. Biervert, et al., A potassium channel mutation in neonatal human epilepsy. Science 279, 403–6 (1998).

27. S. E. Heron, et al., Deletions or duplications in KCNQ2 can cause benign familial neonatal seizures. J. Med. Genet. 44, 791–6 (2007).

28. I. C. Lee, J. J. Yang, S. H. Wong, Y. M. Liou, S. Y. Li, Heteromeric Kv7.2 current changes caused by loss-of-function of KCNQ2 mutations are correlated with long-term neurodevelopmental outcomes. Sci. Rep. 10, 1–14 (2020).

29. C. Hu, D. Liu, T. Luo, Y. Wang, Z. Liu, Phenotypic spectrum and long-term outcome of children with genetic early-infantile-onset developmental and epileptic encephalopathy. Epileptic Disord. 24, 343–352 (2022).

30. N. Dirkx, F. Miceli, M. Taglialatela, S. Weckhuysen, The role of Kv7.2 in neurodevelopment: insights and gaps in our understanding. Front. Physiol. 11, 570588 (2020).

31. E. C. Cooper, E. Harrington, Y. N. Jan, L. Y. Jan, M channel KCNQ2 subunits are localized to key sites for control of neuronal network oscillations and synchronization in mouse brain. J. Neurosci. 21, 9529–40 (2001).

32. E. C. Cooper, et al., Colocalization and coassembly of two human brain M-type potassium channel subunits that are mutated in epilepsy. Proc. Natl. Acad. Sci. U. S. A. 97, 4914–9 (2000).

33. H. S. Wang, et al., KCNQ2 and KCNQ3 potassium channel subunits: molecular correlates of the M-channel. Science 282, 1890–3 (1998).

34. P. Delmas, D. A. Brown, Pathways modulating neural KCNQ/M (Kv7) potassium channels. Nat. Rev. Neurosci. 6, 850–862 (2005).

35. H. Watanabe, et al., Disruption of the epilepsy KCNQ2 gene results in neural hyperexcitability. J. Neurochem. 75, 28–33 (2000).

36. C. Gonzalez, et al., Ribosome profiling reveals a cell-type-specific translational landscape in brain tumors. J. Neurosci. 34, 10924–10936 (2014).

37. S. J. Kiniry, A. M. Michel, P. V. Baranov, The GWIPS-viz browser. Curr. Protoc. Bioinforma. 62, e50 (2018).

38. M. G. Kearse, J. E. Wilusz, Non-AUG translation: a new start for protein synthesis in eukaryotes. Genes Dev. 31, 1717–1731 (2017).

39. C. G. Vanoye, et al., High-throughput evaluation of epilepsy-associated KCNQ2 variants reveals functional and pharmacological heterogeneity. JCI insight 7, 1–14 (2022).

40. Y. Suzuki, et al., An upstream open reading frame and the context of the two AUG codons affect the abundance of mitochondrial and nuclear RNase H1. Mol. Cell. Biol. 30, 5123–34 (2010).

41. T. E. Dever, I. P. Ivanov, M. S. Sachs, Conserved upstream open reading frame nascent peptides that control translation. Annu. Rev. Genet. 54, 237–264 (2020).

42. C. Rabadan-Diehl, A. Martínez, S. Volpi, S. Subburaju, G. Aguilera, Inhibition of vasopressin V1b receptor translation by upstream open reading frames in the 5’-untranslated region. J. Neuroendocrinol. 19, 309–19 (2007).

43. P. Barragan-Iglesias, et al., A peptide encoded within a 5’ untranslated region promotes pain sensitization in mice. Pain 162, 1864–1875 (2021).

44. D. R. Jayaram, et al., Unraveling the hidden role of a uORF-encoded peptide as a kinase inhibitor of PKCs. Proc. Natl. Acad. Sci. U. S. A. 118, 1–11 (2021).

45. H. Yang, Q. Li, E. K. Stroup, S. Wang, Z. Ji, Widespread stable noncanonical peptides identified by integrated analyses of ribosome profiling and ORF features. Nat. Commun. 15, 1932 (2024).

46. S. K. Young, R. C. Wek, Upstream open reading frames differentially regulate gene-specific translation in the integrated stress response. J. Biol. Chem. 291, 16927–35 (2016).

47. C. M. Rodriguez, S. Y. Chun, R. E. Mills, P. K. Todd, Translation of upstream open reading frames in a model of neuronal differentiation. BMC Genomics 20, 1–18 (2019).

48. P. P. Mueller, A. G. Hinnebusch, Multiple upstream AUG codons mediate translational control of GCN4. Cell 45, 201–7 (1986).

49. K. M. Vattem, R. C. Wek, Reinitiation involving upstream ORFs regulates ATF4 mRNA translation in mammalian cells. Proc. Natl. Acad. Sci. U. S. A. 101, 11269–74 (2004).

50. A. G. Hinnebusch, Translational regulation of yeast GCN4. A window on factors that control initiator-trna binding to the ribosome. J. Biol. Chem. 272, 21661–4 (1997).

51. P. K. Stanton, J. M. Sarvey, Blockade of long-term potentiation in rat hippocampal CA1 region by inhibitors of protein synthesis. J. Neurosci. 4, 3080–8 (1984).

52. L. E. Ostroff, et al., Shifting patterns of polyribosome accumulation at synapses over the course of hippocampal long-term potentiation. Hippocampus 28, 416–430 (2018).

53. E. Hacisuleyman, et al., Neuronal activity rapidly reprograms dendritic translation via eIF4G2:uORF binding. Nat. Neurosci. 27, 822–835 (2024).

54. D. J. Kiltschewskij, M. J. Cairns, Transcriptome-wide analysis of interplay between mRNA stability, translation and small RNAs in response to neuronal membrane depolarization. Int. J. Mol. Sci. 21, 1–23 (2020).

55. M. J. Ruiz-Echevarría, S. W. Peltz, The RNA binding protein Pub1 modulates the stability of transcripts containing upstream open reading frames. Cell 101, 741–51 (2000).

56. J. T. Mendell, N. A. Sharifi, J. L. Meyers, F. Martinez-Murillo, H. C. Dietz, Nonsense surveillance regulates expression of diverse classes of mammalian transcripts and mutes genomic noise. Nat. Genet. 36, 1073–8 (2004).

57. J. Zhang, X. Sun, Y. Qian, J. P. LaDuca, L. E. Maquat, At least one intron is required for the nonsense-mediated decay of triosephosphate isomerase mRNA: a possible link between nuclear splicing and cytoplasmic translation. Mol. Cell. Biol. 18, 5272–83 (1998).

58. C. Stockklausner, et al., The uORF-containing thrombopoietin mRNA escapes nonsense-mediated decay (NMD). Nucleic Acids Res. 34, 2355–63 (2006).

59. T. A. Bottorff, H. Park, A. P. Geballe, A. R. Subramaniam, Translational buffering by ribosome stalling in upstream open reading frames. PLoS Genet. 18, e1010460 (2022).

60. L. R. Alagar Boopathy, E. Beadle, A. Garcia-Bueno Rico, M. Vera, Proteostasis regulation through ribosome quality control and no-go-decay. Wiley Interdiscip. Rev. RNA 14, e1809 (2023).

61. A. Raguram, S. Banskota, D. R. Liu, Therapeutic in vivo delivery of gene editing agents. Cell 185, 2806–2827 (2022).

62. P. Nappi, et al., Epileptic channelopathies caused by neuronal Kv7 (KCNQ) channel dysfunction. Pflugers Arch. 472, 881–898 (2020).

63. G. Orhan, et al., Dominant-negative effects of KCNQ2 mutations are associated with epileptic encephalopathy. Ann. Neurol. 75, 382–94 (2014).

64. V. M. Miller, et al., Allele-specific silencing of dominant disease genes. Proc. Natl. Acad. Sci. U. S. A. 100, 7195–200 (2003).

65. S. Aguti, et al., Strategies to improve the design of gapmer antisense oligonucleotide on allele-specific silencing. Mol. Ther. Nucleic Acids 35, 102237 (2024).

66. F. Madeira, et al., The EMBL-EBI Job Dispatcher sequence analysis tools framework in 2024. Nucleic Acids Res. 52, W521–W525 (2024).

67. Q. Li, E. K. Stroup, Z. Ji, Rfoot-seq: Transcriptomic RNase footprinting for mapping stable RNA-protein complexes and rapid ribosome profiling. Curr. Protoc. 3, e761 (2023).

68. Q. Li, H. Yang, E. K. Stroup, H. Wang, Z. Ji, Low-input RNase footprinting for simultaneous quantification of cytosolic and mitochondrial translation. Genome Res. 32, 545–557 (2022).

69. B. J. Joseph, et al., TDP-43-dependent mis-splicing of KCNQ2 triggers intrinsic neuronal hyperexcitability in ALS/FTD. Nat. Neurosci. 28, 2476–2492 (2025).

70. J.-M. DeKeyser, C. H. Thompson, A. L. George, Cryptic prokaryotic promoters explain instability of recombinant neuronal sodium channels in bacteria. J. Biol. Chem. 296, 100298 (2021).

71. C. G. Vanoye, et al., Functional evaluation of human ion channel variants using automated electrophysiology. Methods Enzymol. 654, 383–405 (2021).

72. C. H. Thompson, et al., Epilepsy-associated SCN2A (NaV1.2) variants exhibit diverse and complex functional properties. J. Gen. Physiol. 155, e202313375 (2023).

73. J. Schindelin, et al., Fiji: an open-source platform for biological-image analysis. Nat. Methods 9, 676–82 (2012).

74. K. Clement, et al., CRISPResso2 provides accurate and rapid genome editing sequence analysis. Nat. Biotechnol. 37, 224–226 (2019).

75. G. E. Crooks, G. Hon, J.-M. Chandonia, S. E. Brenner, WebLogo: a sequence logo generator. Genome Res. 14, 1188–90 (2004).

76. R Core Team, R: A language and environment for statistical computing. (2022). Available at: https://www.r-project.org.

77. H. Wickham, ggplot2: elegant graphics for data analysis (Springer-Verlag New York, 2016).

