## Supplemental Information for "An upstream open reading frame represses translation of the neuronal potassium channel KCNQ2"

bioRxiv

### Supplementary Information for
An upstream open reading frame represses translation of the neuronal potassium channel KCNQ2

**Dalton J. Huey,[a] Eduardo Guadarrama,[a] Jean-Marc DeKeyser,[a] Carlos G. Vanoye,[a] Christine Q. Simmons,[a] Qianru Li,[a] Emily K. Stroup,[a] Zhe Ji,[a] and Alfred L. George, Jr.[a,*]**
[a]Department of Pharmacology, Feinberg School of Medicine, Northwestern University, Chicago, IL, USA 60611

**This PDF file includes:**

Figures S1 to S4
Tables S1 to S3

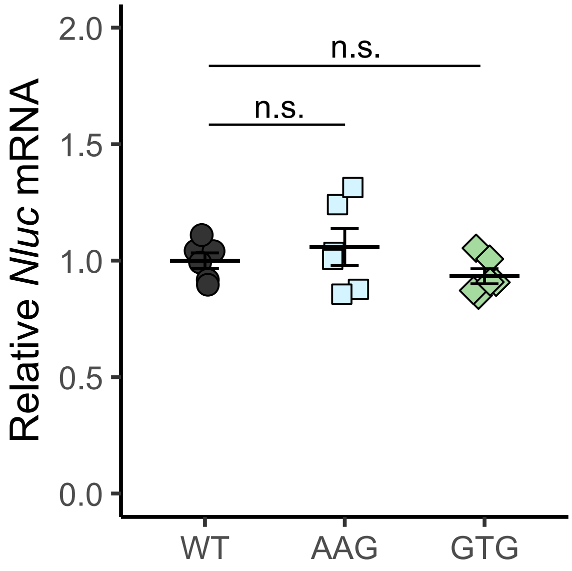


**Fig. S1:** Relative luciferase mRNA levels for *Nluc* in HEK293 cells transiently transfected with reporter gene constructs.

The nanoluciferase CDS was preceded by the *KCNQ2* 5'-UTR containing either the wildtype (WT, dark gray circles), AAG mutated (blue squares), or GTG mutated (green diamonds) uORF sequence (Games-Howell post-hoc, n.s. $p>0.05$, mean ± SEM, n=2 per three transfection replicates, n=6 per condition). *Nluc* mRNA levels are relative to *Fluc* mRNA from co-transfected firefly luciferase constructs as a transfection efficiency control.

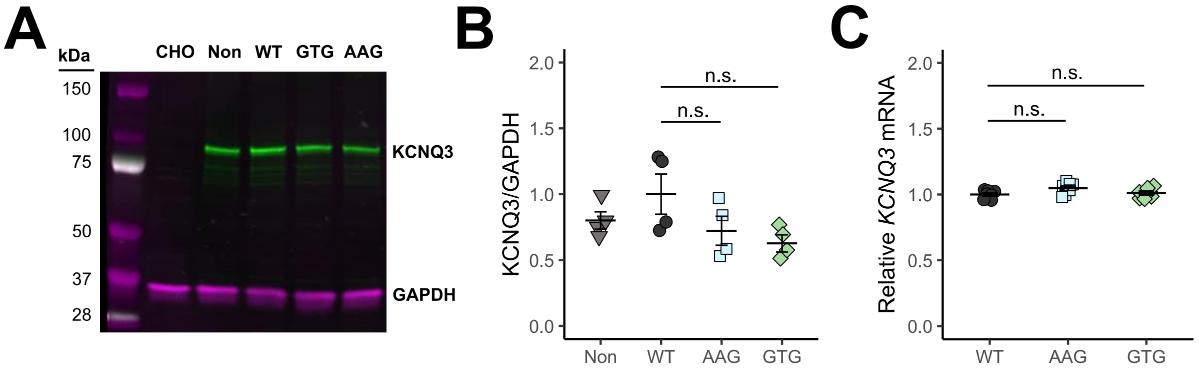


**Fig. S2:** *KCNQ3* protein and mRNA levels are unaffected by uORF mutations

**(A)** Representative western blot for KCNQ3 (green) and GAPDH (magenta) for CHO-Q3 cells electroporated with full length *KCNQ2* constructs. CHO cells lacking *KCNQ3* stable expression were included as a negative control. **(B)** Quantification of KCNQ3 protein signal normalized to GAPDH, relative to the WT condition (Games-Howell post-hoc, n.s. $p>0.05$, mean ± SEM, $n=4$) **(C)** Relative mRNA levels for *KCNQ3* from CHO-Q3 cells electroporated with *KCNQ2* constructs (Games-Howell post-hoc, n.s. $p>0.05$, mean ± 95% CI, $n=9$). Legend: nontransfected cells (light gray triangles), wildtype uORF (WT, dark gray circles), AAG mutation (blue squares), and GTG mutation (green diamonds).

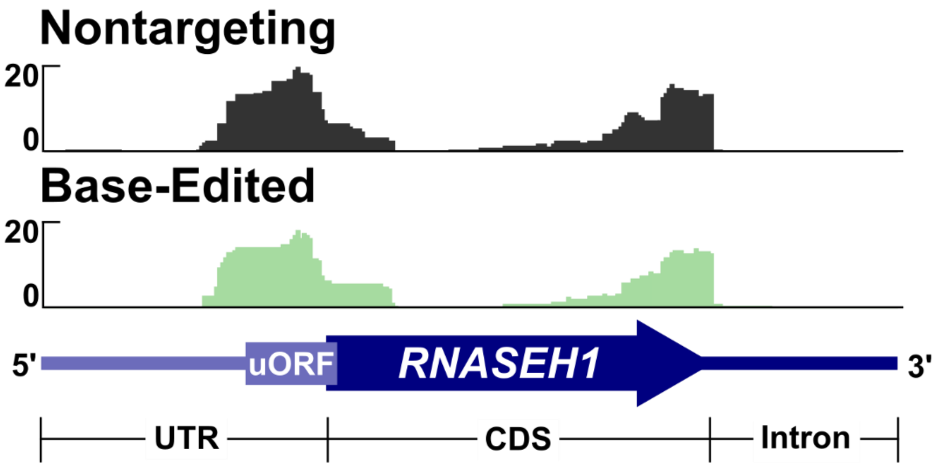


**Fig. S3:** *KCNQ2* uORF base-editing does not alter uORF translation of an unrelated gene

Ribosome profiling of the NT (dark gray) and GTG uORF-edited (light green) SH-SY5Y cell lines for the *RNASEH1* 5'-UTR and 5'-proximal CDS region. Ribosome footprints are shown as combined signal from three independently electroporated cell lines per condition.

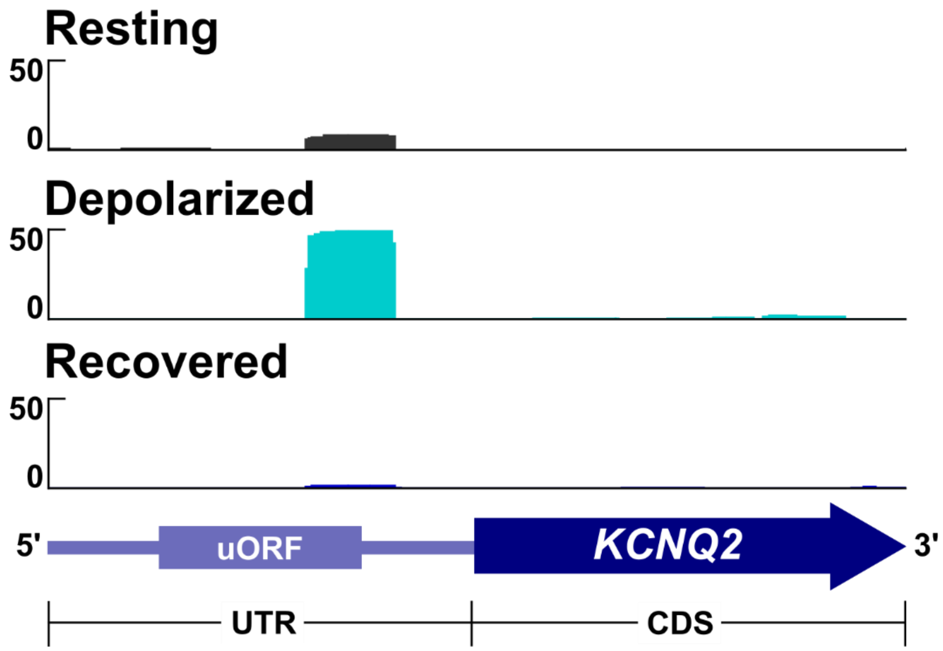


**Fig. S4:** *KCNQ2* uORF translation is enhanced after repeated excitation in a neuron-like cell model

Ribosome profiling of *KCNQ2* from published datasets on differentiated SH-SY5Y cells at rest (dark gray), immediately following repeated depolarization (cyan), and after a 2-hour recovery period (dark blue). Mapped reads of ribosome-protected footprints are shown for the uORF and the 5'-proximal *KCNQ2* CDS. Reads are combined from four independent samples per condition.

#### **Table S1:** Ribo-seq library preparation primers

| Purpose | Sequence |
|---|---|
| SMART-RT primer | ACGTGTGCTCTTCCGATCTNNNNNNNNNNNNNNNTTTTTTTTTTTTTTTVN |
| SMARTer oligo | CTCTTTCCCTACACGACGCTCTTCCGATCTNNNNrGrG+G (rGrG+G locked nucleotides) |
| PCR1 forward primer | GATCTACACTCTTTCCCTACACGACGC |
| PCR 1 reverse primer | GTGACTGGAGTTCAGACGTGTGCTCTTCCGATCT |
| PCR 2 forward primer | AATGATACGGCGACCACCGAGATCTACACTCTTTCCCTACAC |
| PCR 2 reverse primer | CAAGCAGAAGACGGCATACGAGATNNNNNNGTGACTGGAGTTCAGACGTGTG (6nt indices) |

#### **Table S2**: Primers used for cloning and mutagenesis

| Purpose | Forward Primer | Reverse Primer |
|---|---|---|
| Human *KCNQ2* 5'UTR NLuc Insert | GTTTAGTGAACCGTCAGATCACTAGAAGCTT**AGGCGCCGAGGTGCGCGCGG** | CGAAATCTTCGAGTGTGAAGACCATGGT**GCCTGGCGGGAGGCGCCC** |
| Mouse *Kcnq2* 5'UTR NLuc Insert | GTTTAGTGAACCGTCAGATCACTAGAAGCTT**GAACGCAGAGCCGCAGCCGC** | CGAAATCTTCGAGTGTGAAGACCATGGT**GCCTGCGGCGCCCTATC** |
| CMV-NLuc Vector | **ATGGTCTTCACACTCGAAGATTTCG** | **AAGCTTCTAGTGATCTGACGGTTCACTAAAC** |
| *KCNQ2* 5'UTR-CDS Insert | TGCCTGGGGAC**AGGCGCCGAGGTGCGCGCGGA** | ACTTCTGCACCAT**GGTGCCTGGCGGGAGGCGCCC** |
| *KCNQ2* 5'UTR-CDS Vector | GCCAGGCACC**ATGGTGCAGAAGTCGCGCAA** | CTCGGCGCCT**GTCCCCAGGCAGAATGGCGGTT** |
| Human uORF GTG | CCTCGGCC**G**TGCGGCTCCCGGCCGGGG | GGAGCCGCA**C**GGCCGAGGCGGCGGTTCCGCA |
| Human uORF AAG | CTCGGCCA**A**GCGGCTCCCGGCCGGGGG | GGGAGCCGC**T**TGGCCGAGGCGGCGGTTCCGC |
| Mouse uORF AAG | CTCGCCA**A**GCGGCTGCCGGACCGGAGG | CCGGCAGCCGC**T**TGGCGAGGCGGCTGCGG |
| uORF Kozak -3G:C | CCGCCTCG**C**CCATGCGGCTCCCGGCCG | GCCGCATGG**G**CGAGGCGGCGGTTCCGCACTC |
| uORF Kozak -2C:G | CGCCTCGG**G**CATGCGGCTCCCGGCCGG | AGCCGCATG**C**CCGAGGCGGCGGTTCCGCACT |
| uORF Kozak -1C:G | GCCTCGGC**G**ATGCGGCTCCCGGCCGGGG | GAGCCGCAT**C**GCCGAGGCGGCGGTTCCGCAC |
| Human uORF TTG | CCTCGGCC**T**TGCGGCTCCCGGCCGGGG | GGAGCCGCA**A**GGCCGAGGCGGCGGTTCCGCA |
| Human uORF ATC | TCGGCCAT**C**CGGCTCCCGGCCGGGGG | CGGGAGCCG**G**ATGGCCGAGGCGGCGGTTCCG |

#### **Table S3**: Custom Taqman Assays

| CHO *B2M* Taqman Assay | Sequence |
|---|---|
| Forward Primer | CCCACTGCGACCGATAAATA |
| Reverse Primer | CAGACCTCCATGATGCTTGA |
| Probe | CACACCACTCTGAAGGAGCCCAAG (HEX fluorophore) |
| *Fluc* Luciferase Taqman Assay | Sequence |
| Forward Primer | CTTCGAGGAGGAGCTATTCTTG |
| Reverse Primer | GTCGTACTTGTCGATGAGAGTG |
| Probe | TGCTGGTGCCCACACTATTTAGCT (Cy5 fluorophore) |
| *Nluc* Luciferase Taqman Assay | Sequence |
| Forward Primer | TTTAAGGTGGTGTACCCTGTG |
| Reverse Primer | CATACGGCCGTCCGAAATA |
| Probe | AAGGTGATCCTGCACTATGGCACA (6-FAM fluorophore) |
